# Multi-model Order ICA: A Data-driven Method for Evaluating Brain Functional Network Connectivity Within and Between Multiple Spatial Scales

**DOI:** 10.1101/2021.10.24.465635

**Authors:** Xing Meng, Armin Iraji, Zening Fu, Peter Kochunov, Aysenil Belger, Judy M. Ford, Sara McEwen, Daniel H. Mathalon, Bryon A. Mueller, Godfrey Pearlson, Steven G. Potkin, Adrian Preda, Jessica Turner, Theo G.M. van Erp, Jing Sui, Vince D. Calhoun

## Abstract

**Background:** While functional connectivity is widely studied, there has been little work studying functional connectivity at different spatial scales. Likewise, the relationship of functional connectivity *between* spatial scales is unknown.

**Methods:** We proposed an independent component analysis (ICA) - based approach to capture information at multiple model orders (component numbers) and to evaluate functional network connectivity (FNC) both within and between model orders. We evaluated the approach by studying group differences in the context of a study of resting fMRI (rsfMRI) data collected from schizophrenia (SZ) individuals and healthy controls (HC). The predictive ability of FNC at multiple spatial scales was assessed using support vector machine (SVM)-based classification.

**Results:** In addition to consistent predictive patterns at both multiple-model orders and single model orders, unique predictive information was seen at multiple-model orders and in the interaction between model orders. We observed that the FNC between model order 25 and 50 maintained the highest predictive information between HC and SZ. Results highlighted the predictive ability of the somatomotor and visual domains both within and between model orders compared to other functional domains. Also, subcortical-somatomotor, temporal-somatomotor, and temporal-subcortical FNCs had relatively high weights in predicting SZ.

**Conclusions:** In sum, multi-model order ICA provides a more comprehensive way to study FNC, produces meaningful and interesting results which are applicable to future studies. We shared the spatial templates from this work at different model orders to provide a reference for the community, which can be leveraged in regression-based or fully automated (spatially constrained) ICA approaches.

**Impact Statement:** Multi-model order ICA provides a comprehensive way to study brain functional network connectivity within and between multiple spatial scales, highlighting findings that would have been ignored in single model order analysis. This work expands upon and adds to the relatively new literature on resting fMRI-based classification and prediction. Results highlighted the differentiating power of specific intrinsic connectivity networks on classifying brain disorders of schizophrenia patients and healthy participants, at different spatial scales. The spatial templates from this work provide a reference for the community, which can be leveraged in regression-based or fully automated ICA approaches.

## Introduction

Brain activity reveals exquisite coordination between spatial scales, from local microcircuits to brain-wide networks. Brain activity measured by noninvasive functional brain imaging techniques is typically assumed to be generated on the cortical surface. The spatial extent of activity on the cortex obtained experimentally from neuroimaging modalities and models vary widely (Perdue and Diamond 2013). Understanding the brain requires an integrated understanding of the different scales of the spatial organization of the brain. Studies that utilize measurements between spatial scales promise to increase our understanding of brain function by tracking sensory, motor, and cognitive variables as they evolve through local microcircuits and across brain-wide networks (Lewis, Bosman, and Fries 2015).

Whole-brain functional connectivity can be studied by calculating the functional network connectivity (FNC) (Allen et al. 2011) between intrinsic connectivity networks (ICNs), *i*.*e*., the fMRI independent components (IC) resulted from group independent component analysis (ICA). To date, most previous research has focused on FNC within one specific model order (*i*.*e*., number of components), ignoring the importance of capturing functional information at different levels of spatial granularity as well as the between-order information. A wide number of studies have focused on FNC in brain health and various disorders, all at a single model order. In particular, multiple studies have highlighted significant FNC differences in studies of schizophrenia (SZ), e.g., increased FNC in SZ among frontal and temporal networks (Vince D. Calhoun, Eichele, and Pearlson 2009), within the default mode networks (Whitfield-Gabrieli et al. 2009; Salvador et al. 2010), and decreased FNC between temporal and parietal networks (Jafri MJ et al. 2008). While it is well-known that there are functional changes associated with schizophrenia, it is unclear to what degree these are linked to the choice of a specific model order (IC numbers) when performing FNC analysis.

Considering the brain as a functional hierarchy (Iraji et al. 2019 a), it is clear that functional interactions between functional sources can occur between different spatial scales, and the lack of studies that evaluate functional connectivity between spatial scales represents a gap in the field. Here, we propose to study FNC within and between four different functional hierarchy levels (model order = 25, 50, 75, and 100), thus providing new insight to understand brain functional connectivity. In this study, our goal is to compare the important information obtained at different model orders by evaluating the multi-order FNCs using HC-SZ group comparison, as well as testing the classification power of FNC in discriminating SZs based on machine learning-based approaches (Pariyadath, Stein, and Ross 2014; Anderson and Cohen 2013). We seek to identify the FNC-based features that are predictive of schizophrenia from controls through SVM-based classifications. Specifically, we evaluated if the HC-SZ group differences primarily occur at one specific model order or the between-model orders, and which order yields the highest classification power between HC and SZ. The ICNs template at four model orders was also made available to the community for their future use.

## Methods

### Dataset and preprocessing

Our dataset was combined across three separate studies, one with seven sites (FBIRN: Functional Imaging Biomedical Informatics Research Network), one with three sites (MPRC: Maryland Psychiatric Research Center), and one single site (COBRE: Center for Biomedical Research Excellence). We extracted a subset of subjects from the three datasets that satisfying the following criteria, 1) data of individuals with typical control or schizophrenia diagnosis; 2) data with high-quality registration to echo-planar imaging (EPI) template; 3) head motion transition of less than 3° rotations and 3-mm translations in all directions. The mean framewise displacement (meanFD) among selected subjects was average ± standard deviation = 0.1778 ± 0.1228; min ∼ man = 0.0355 ∼ 0.9441. This resulted in 827 individuals, including 477 subjects (age: 38.76 ± 13.39, females: 213, males: 264) of HCs and 350 SZ individuals (age: 38.70 ± 13.14, females: 96, males: 254). The parameters of resting-state fMRI (rsfMRI) for the FBIRN data were the same across all sites, with a standard gradient echo-planar imaging (EPI) sequence, repetition time (TR)/echo time (TE) = 2000/30 ms, and a total of 162 volumes. For COBRE data, rsfMRI images were acquired using a standard EPI sequence with TR/TE = 2000/29 ms and 149 volumes. The MPRC dataset were acquired using a standard EPI sequence in three sites, including Siemens 3-Tesla Siemens Allegra scanner (TR/TE = 2000/27 ms, voxel spacing size = 3.44 × 3.44 × 4 mm, FOV = 220 × 220 mm, and 150 volumes), 3-Tesla Siemens Trio scanner (TR/TE = 2210/30 ms, voxel spacing size = 3.44 × 3.44 × 4 mm, FOV = 220 × 220 mm, and 140 volumes), and 3-Tesla Siemens Tim Trio scanner (TR/TE = 2000/30 ms, voxel spacing size = 1.72 × 1.72 × 4 mm, FOV = 220 × 220 mm, and 444 volumes).

We were following the same rsfMRI data preprocessing procedures as (A. Iraji et al. 2021) did. First of all, for the purposes of magnetization equilibrium, the first five volumes were discarded. We performed rigid motion correction to correct the head motion for each subject during scanning. We then applied slice-time correction to deal with temporal misalignment. And the next step, the rsfMRI data of each subject was registered to a Montreal Neurological Institute (MNI) EPI template, resampled to 3 mm^3^ isotropic voxels using, and smoothed using a Gaussian kernel with a 6 mm full width at half-maximum (FWHM = 6 mm). The voxel time courses were then z-scored (variance normalized). The preprocessing procedures were mainly performed by using the statistical parametric mapping (SPM12, https://www.fil.ion.ucl.ac.uk/spm/) toolbox.

### Group ICA analysis

ICA provides adaptive, overlapping, networks at different scales, we and others have shown this has substantial advantages over fixed ROIs (Y. Du et al. 2020; Yu et al. 2018). The main advantages including the ability to adapt to individual subjects, the fact that the maps are optimized to include temporally coherent voxels, the accounting for overlapping networks, which we have shown captures important information, and the continuous nature of the maps, providing a more natural decomposition of the data. In ICA analysis, the selection of model order (i.e., number of components to be extracted) can effectively define spatial scale of ICNs and therefore has crucial effects on brain functional network analysis. This makes ICA a great tool to obtain subject-specific ICNs at different spatial scales. In another word, we can study brain functionwithin and between different spatial scales by using ICA with different model orders. Low model order ICAs produce large-scale spatially distributed ICNs, such as the default mode network (Beckmann et al. 2005; Damoiseaux et al. 2008; Vince D. Calhoun, Kiehl, and Pearlson 2008), whereas high model orders produce spatially granular ICNs, including multiple fine ICNs instead of one large-scale the default mode network (Allen et al. 2011). Given these advantages, we tend to study brain function at multiple scales using ICA.

The ICA analysis was performed using the GIFT software (http://trendscenter.org/software/gift, Vince D. Calhoun and Adali 2012; Vince D. Calhoun, Adali, et al. 2001). In our study, ICA was performed at four model orders: 25, 50, 75, and 100. Prior to the ICA, subject-specific principal components analysis (PCA) was first performed on the datasets to normalize the data. The subject-level principal components were then concatenated together across the time dimensions, and the group-level PCA was applied to concatenate the subject-level principal components. The group-level principal components that explained the maximum variance were selected as the input of ICA to perfrom group-level ICA (Vince D. Calhoun et al. 2001). We used the infomax algorithm and controlled for stochastic variability by using ICASSO (Himberg, Hyvärinen, and Esposito 2004) as implemented in the GIFT software by running ICA several times and selecting the most representative run (W. Du et al. 2014). ICASSO was used to evaluate component stability. The Infomax ICA algorithm was run for 100 times and clustered together within the ICASSO framework (Himberg, Hyvärinen, and Esposito 2004). The run with the closest independent components to the centrotypes of stable clusters (ICASSO cluster quality index > 0.8) (Armin Iraji et al. 2020) was selected as the best run and used for future analysis (Ma et al. 2011).

We utilized ICA with different model orders (25, 50, 75, and 100) to identify ICNs at multiple spatial scales. ICNs were identified from each model order and included components with peak activations in gray matter and low-frequency timecourses (Vince D. Calhoun, Eichele, and Pearlson 2009). The subject-specific ICNs time courses were calculated using the spatial multiple regression technique (Vince D. Calhoun, Pekar, and Pearlson 2004). Prior to calculating static FNC, an additional cleaning procedure was performed on the time courses of ICNs to reduce the effect of the remaining noise and improve the detection of FNC patterns. The procedures were as follows: 1) Remove the linear, quadratic, and cubic trends, 2) Regress out the motion realignment variables 3) Replace outliers with the best estimate using a third-order spline fit, and 4) Bandpass filter using a fifth-order Butterworth filter with frequency cutoff of 0.01 Hz-0.15 Hz (Allen et al. 2011).

### FNC analysis at multiple spatial scales

FNC was computed between each pair of ICNs by calculating the Pearson correlation coefficient between ICNs (V D Calhoun et al. 2003; Allen et al. 2011; Madiha J Jafri et al. 2008). We calculated a 2D symmetric ICN × ICN FNC matrix for each subject and aggregated the FNC matrix from all subjects into an augmented 2D matrix. We then calculate the mean FNC matrix of all subjects for further analysis.

To investigate the group differences between HC and SZ in FNC, we performed a generalized linear model (GLM). We fit the GLM model with age, gender, data acquisition site, and meanFD as covariates. The meanFD was added to the GLM to account for any residual motion effect that was not removed in the preprocessing steps. The group differences between HC and SZ in FNC were evaluated by the t-value and p-value of statistical comparisons of the GLM.

### Support vector machine-based classification (SVM)

The SVM (Verma and Salour, Al 1992) is so far the most popular classification method due to its favorable characteristics of high accuracy, ability to deal with high-dimensional data, and versatility in modeling diverse sources of data. The SVM has been widely applied in numerous neuroimaging classification studies and has achieved remarkable results due to its excellent generalization performance. Our motivation of using SVM over other approaches was due to its sensitivity, resilience to overfitting, ability to extract and interpret features, and superior performance in fMRI data classification (Wang et al. 2019; De Martino et al. 2008; Pereira, Mitchell, and Botvinick 2009; Ecker et al. 2010; Liu et al. 2013; Vergun et al. 2013; Saha et al. 2021). To investigate the group differences, we applied a binary SVM nonlinear Gaussian RBF kernel classifier (Chih-Wei Hsu, Chih-Chung Chang, and Chih-Jen Lin 2003), due to the fact that the Gaussian RBF kernel generally outperforms other kernels in brain MRI imaging classification (Madheswaran and Anto Sahaya Dhas 2015; Kumari 2013). The classification model was trained and cross-validated using the dataset of 477 HCs and 350 SZs. The figure (Fig.1) presents the pipeline for training and testing the SVM model.

**Fig. 1.**
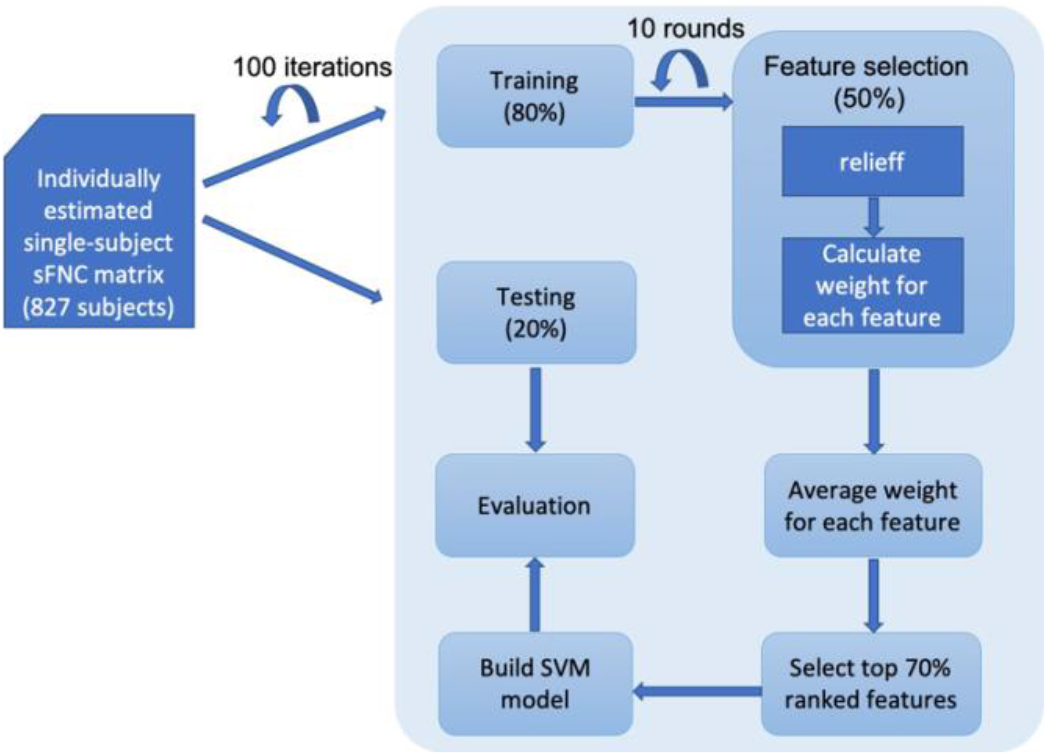
Pipeline of the classification model. The whole dataset was split into a training set (80%) and a testing set (20%). Feature selection was performed on the training set (50% were selected randomly every time). In order to select the most predictive features, we repeated the feature selection process for ten rounds and retained those features with a high average weight (top 70%) among all the rounds. The final SVM model was built based on the selected FNC features. We ran the modeling process for a total number of 100 iterations to obtain a stable SVM model.

### Feature selection

Each FNC pair of the functional connectivity among ICNs was considered as the input feature for classification, and the category of group HC or SZ was considered as the response vector. Given that some of those FNC features might be non informative or redundant for classification, we performed feature selection using Relief (Verma and Salour, Al 1992) to improve classification performance and to speed up computation. Relief ranks predictors with *k* nearest neighbors (we set *k* to 10 for calculation simplicity). The function returns the indices of the most important predictors and the weights of the predictors. Feature selection was carried out prior to classifier training through the recursive feature elimination step. For each round of feature selection, 50% of the training data (as shown in Fig. 1) was selected. We repeated it for ten rounds and retained those features with a high average weight (top 70%) among all the rounds. We thus narrowed the set of features to a subset of the original feature set, which eliminated the non-informative and redundant FNC features. The SVM model was built based on the selected FNC feature set.

### Recursive validation

In order to obtain a stable performance of the SVM model, we applied recursive validation. For each iteration, we randomly split the whole dataset into 80% of the training set and 20% of the testing set. The test set was held out for final evaluation. We ran the modeling process for a total number of 100 iterations and evaluated the SVM model based on average specificity, sensitivity, and F1 score (the harmonic mean of the precision and recall) across all iterations.

## Results

### ICA analysis

Spatial maps of selected ICs at different model orders are shown in the figure (Fig. 2 a – d). A total of 127 ICNs were determined between all model orders (15, 28, 36, and 48 from 25, 50, 75, and 100, respectively). ICNs were grouped into functional domains including cerebellum, cognitive control, default mode, somatomotor, subcortical, temporal, and visual (Fig. 3).

**Fig. 2.**
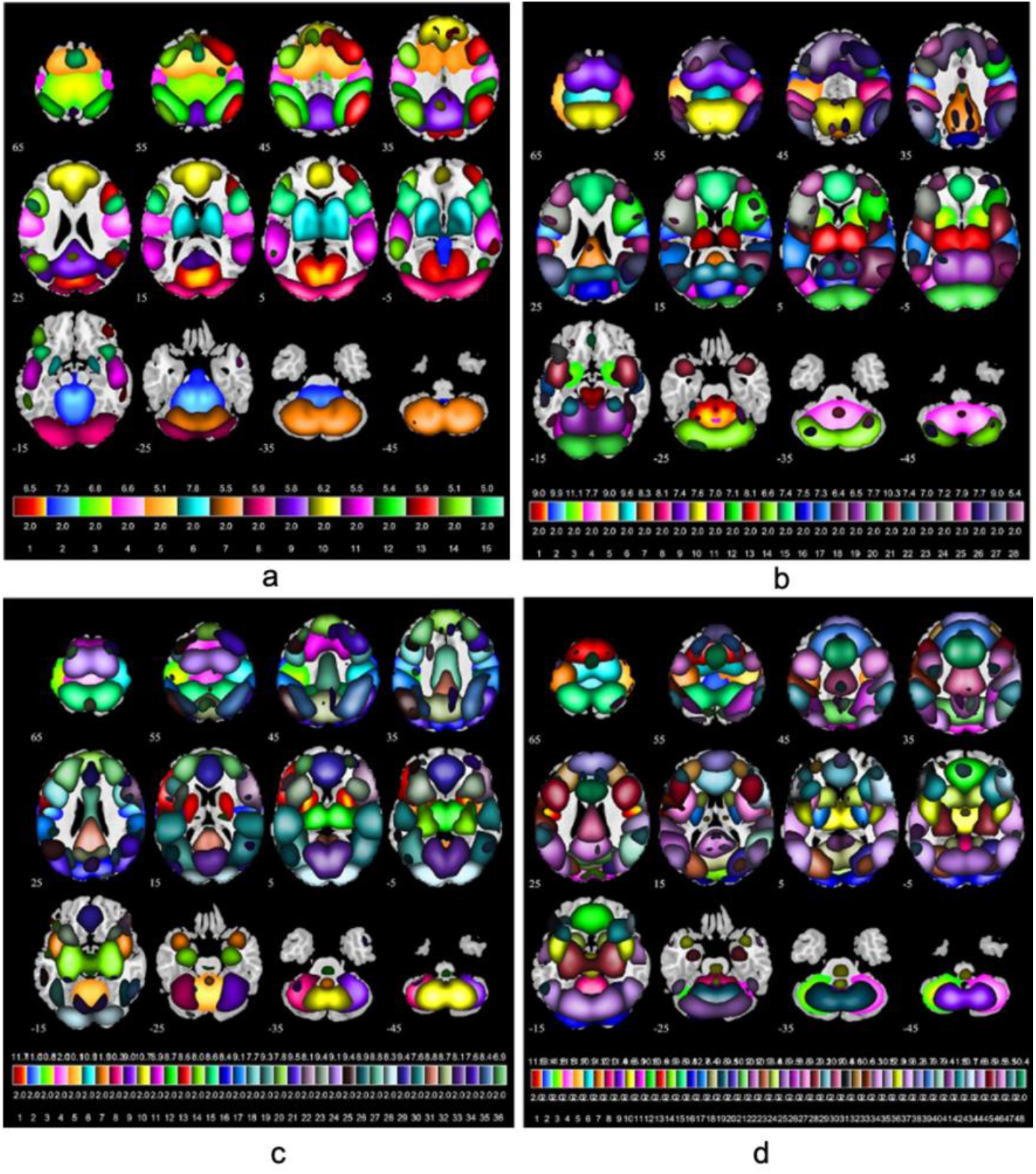
ICN maps selected from model order of 25 (2 a), 50 (2 b), 75 (2 c), and 100 (2 d). A total number of 127 ICNs were determined between all model orders (15, 28, 36, and 48 from 25, 50, 75, and 100, respectively). All the ICNs were identified from each model order and included components with peak activations in gray matter as well as low-frequency time-courses.

**Fig. 3.**
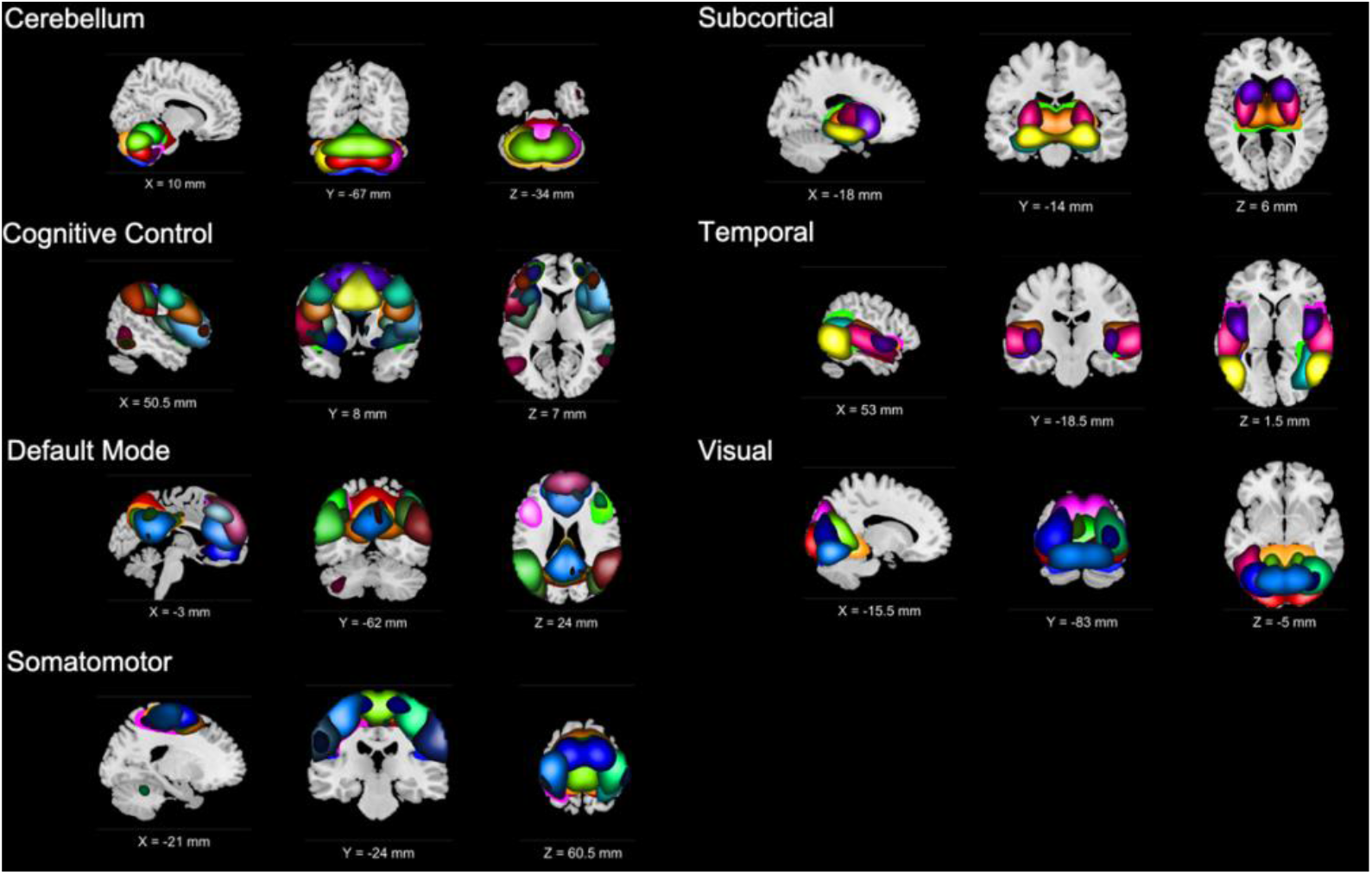
Identified ICNs across all model orders 25, 50, 75, and 100. ICNs were divided into groups (functional domains) based on their anatomical and functional properties and include cerebellum (CR), cognitive control (CC), default mode (DM), somatomotor (SM), subcortical (SB), temporal (TP), and visual (VS). Each functional domain is displayed at the three most informative slices.

The results show that at lower model orders (25 ∼ 50) signal sources tend to merge into singular components, which then split into several subcomponents at higher model orders. These findings are consistent with previous and most recent findings (Abou-Elseoud et al. 2010; Rachakonda, Du, and V. D. Calhoun 2017). Figures (Fig. 2 c – d) show that higher model order ICA (75 ∼ 100) tends to parcellate the brain into focal functionally homogeneous distinct regions, which are consistent with previous studies (Iraji et al. 2009 b; Calhoun, VD and T 2012; V. Calhoun and N. 2017). The ICN templates (we show in Fig. 2 and Fig. 3) are openly shared at http://trendscenter.org/data.

### FNC analysis

The figures (Fig. 4 a – b) present the mean FNC (z-fisher score) between 127 ICNs between all model orders. The figure (Fig. 4 a) was sorted by ICNs and then model orders within each domain (called an FNC ‘block plot’), and the figure (Fig. 4 b) was sorted by model order and then the domain (called an FNC ‘finger plot’), aiming to visualize patterns existing at and between different model orders. In the block plot, we observed that the cerebellum domain was highly correlated with itself, anticorrelated across all but subcortical domains. The somatomotor, subcortical, visual (and to a lesser degree temporal) domains were more homogeneous than the default mode, cerebellum, and cognitive control domains both within and between model orders. We also observed strong anticorrelations between default mode vs. somatomotor and temporal between model order 25 and 100 in the finger plot, which were not clearly seen in the other model orders.

**Fig. 4.**
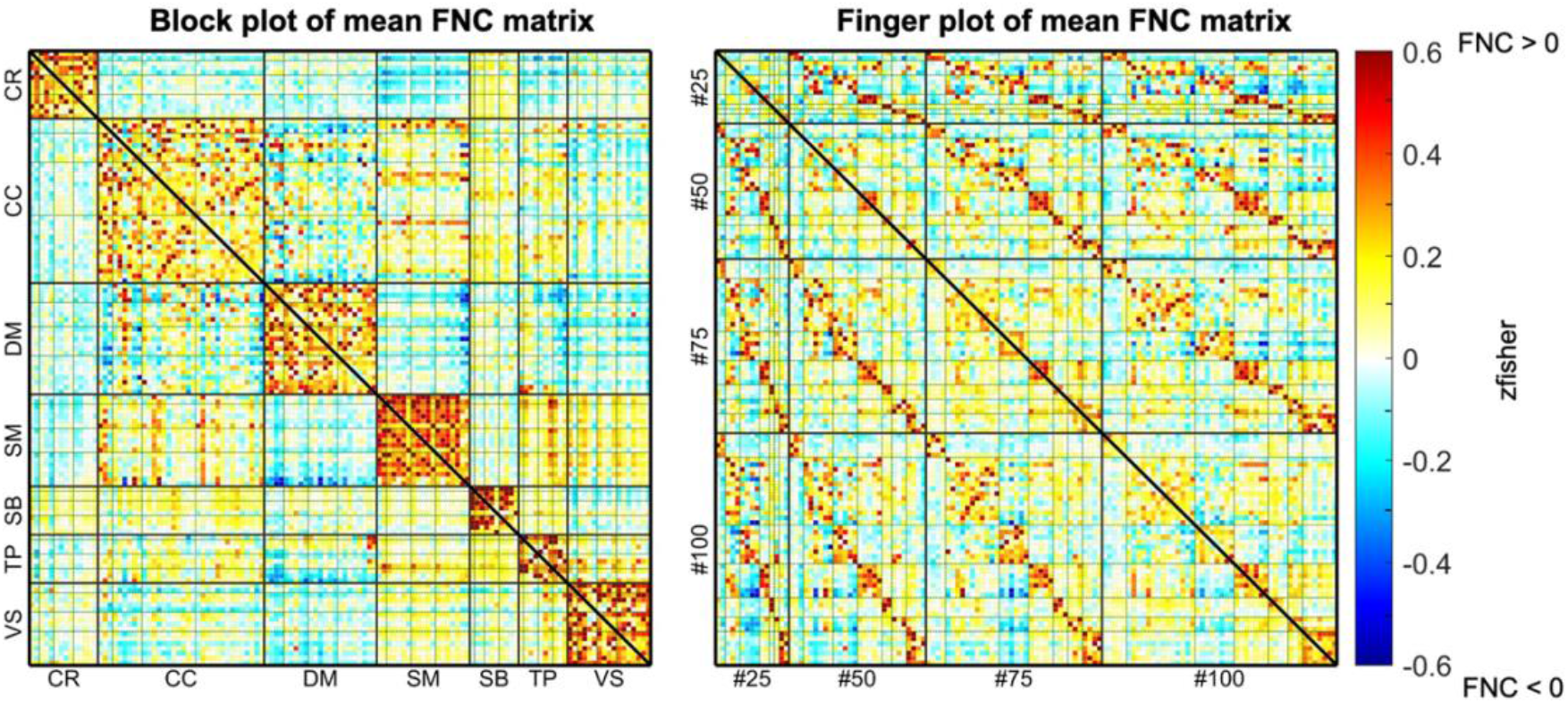
a (left). Average FNC plots. We calculated the mean FNC (z-fisher score) based on the aggregated FNC matrix of all individuals. Fig. 4 a (left). Block plot of mean FNC matrix between model orders. The ICNs in this FNC matrix were sorted by domains first, and within each domain, ICNs were sorted by model orders (from 25 to 100). The dot lines in each domain divide different model orders. Fig. 4 b (right). Finger plot of mean FNC matrix between model orders. ICNs within each model order were sorted in the order of cerebellum (CR), cognitive control (CC), default mode (DM), somatomotor (SM), subcortical (SB), temporal (TP), and visual (VS). As we can see in the plot, DM from model order 25 show strong anticorrelations with SM and TP of model order 100; similar anticorrelations were also observed between model order 75 and 100, which did not show up in the other sections of the matrix.

We also evaluated the group differences between schizophrenia and controls (Fig. 5 a - b). Strong increases were observed between the ICNs of the subcortical domain and the ICNs of the temporal, visual, and somatomotor domains in the SZ group. A similar increased FNC in SZ was observed between the ICNs of the cerebellum domain and the ICNs of somatomotor, temporal, and visual domains. The most dominant decreases in FNC in the SZ group were found mainly within the ICNs of the somatomotor domain and between the ICNs of the somatomotor domain and the ICNs of the temporal and visual domains. Furthermore, slight decreases in SZ were seen between the cerebellum and subcortical compared to the HC group.

**Fig. 5.**
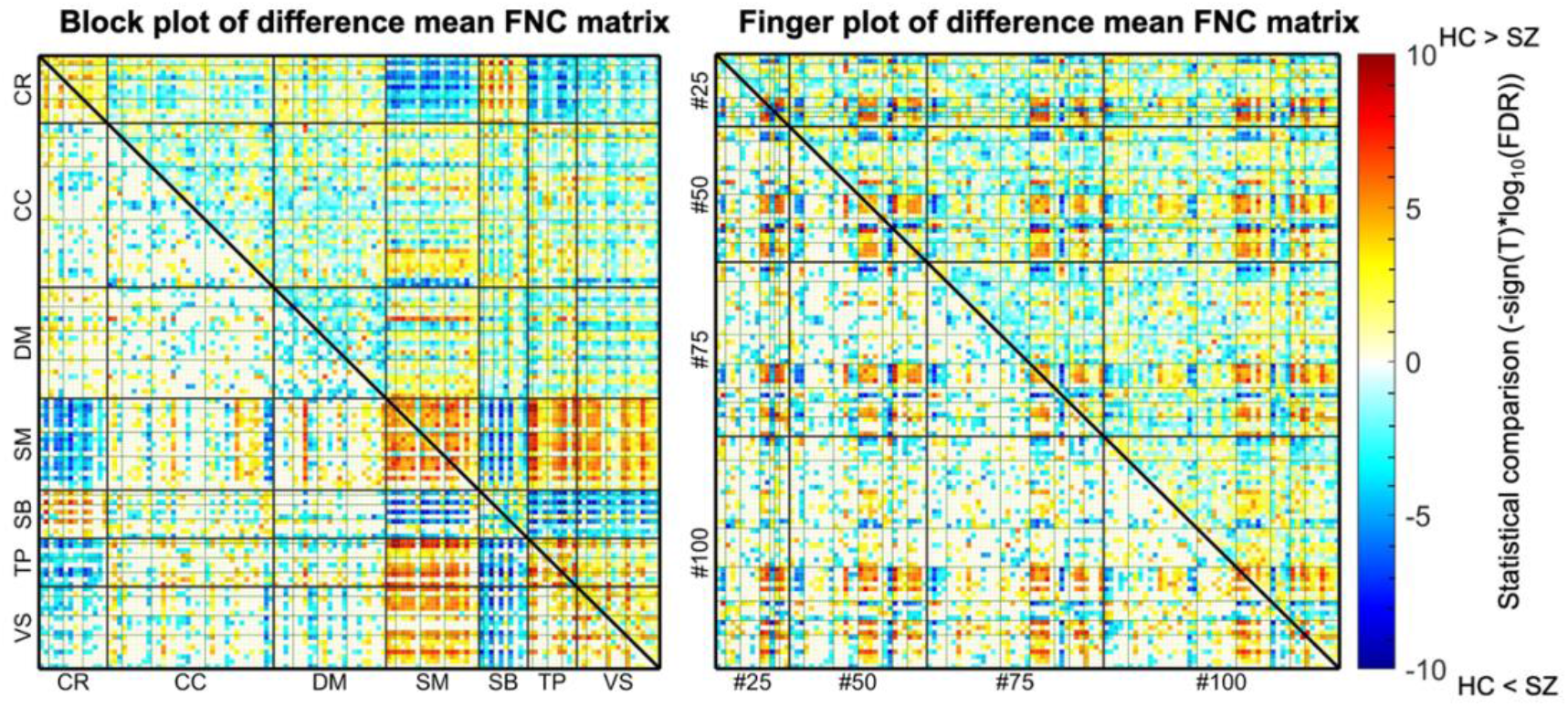
a (left). Block plot of GLM contrast of difference mean FNC matrix (HC - SZ). To test the group difference between HC and SZ, we fit a GLM with age, gender, data acquisition site, and meanFD as covariates. This figure shows the difference between HC and SZ in FNC. Blue areas indicate increased FNC in SZ compared to HC, and red areas indicate decreased FNC in SZ compared to HC. The statistical results of p values of the GLM were corrected for multiple comparisons using a 5% false discovery rate (FDR). Fig. 5 b (right). Finger plot of generalized linear model contrast of difference mean FNC matrix (HC - SZ). Fig. 5 a - b show the intensity (-sign(T) ^*^log10(FDR)) matrixes of both block and finger plots for the mean FNC of the GLM model, where T is the t-statistic values of the GLM. The lower triangles show covariate pairs that were significantly different (FDR < 0.05).

### SVM model

The figure (Fig. 6 a) and b represent the block plot and the finger plot of average weights of FNC features of 100 iterations. This gives us a better view of the importance of each FNC feature in detecting the group differences. As shown in the figures (Fig. 6 a - b), FNC features in the cerebellum, somatomotor, subcortical, temporal, and visual domains consistently contribute to the classification results for both within and between different model orders. These findings suggest highly distinct functional roles for these ICNs in differentiating between HCs and SZs. It is noticeable that the FNC within the somatomotor and visual have contributed most to the classification. The predictive abilities of the FNC features within those two domains were evenly distributed at all model orders, as they are shown in the diagonal regions within somatomotor and visual (in Fig. 6 a). Generally, the strong predictive abilities of FNC features were observed in the FNC pairs between the subcortical domain and the domains of the cerebellum, the somatomotor, and the temporal across all model orders. Similarly, FNC features between the somatomotor and the domains in the cerebellum and temporal were generally predictive.

**Fig. 6.**
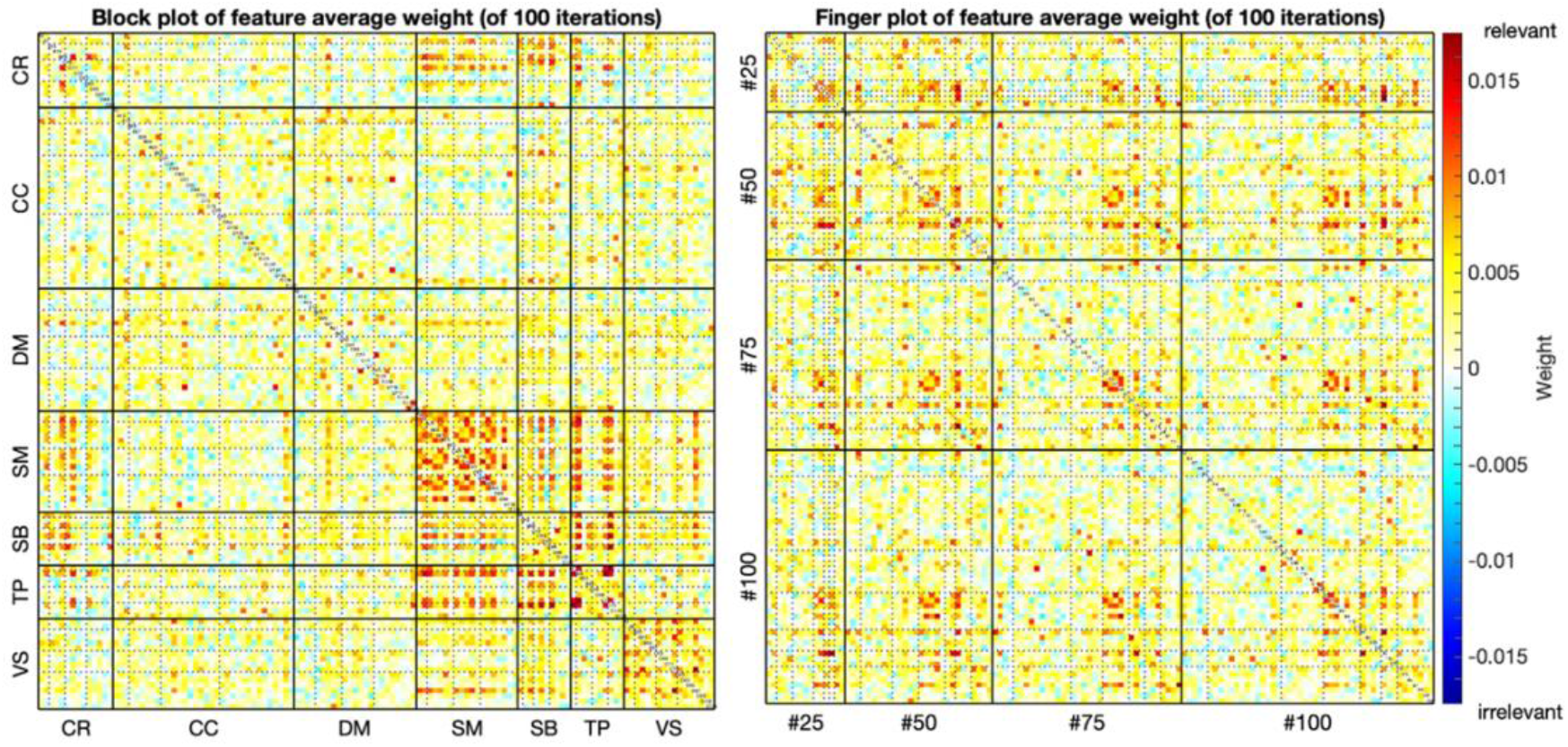
a (left). SVM block plot of FNC feature average weights of 100 iterations. The hotspots indicate FNC features with strong weights in predicting group differences of HC and SZ. Statistically, the relevance level of a relevant feature is expected to be larger than zero and that of an irrelevant one is expected to be zero (or negative). The cerebellum, somatomotor, subcortical, and temporal domain almost always contribute to the classification. Generally, strong predictive abilities of FNC features in the somatomotor and visual domain were seen at all model orders. Visual was predictive only within the region between somatomotor and visual, but only seen at high model order (100). Fig. 6 b (right). SVM finger plot of FNC feature average weights of 100 iterations.

To better understand the between model order differences, we computed a two-sample t-test (Cressie and Whitford 1986), to identify significant differences within and between model orders. For each iteration, we calculated an average feature weight for each model order, we then aggregated all the average feature weights across 100 iterations. The two-sample t-tests were performed between all pairs of model orders using their aggregated feature weights. It (Fig. 7) shows the statistical comparison (intensity) plot between model orders. A negative intensity value in the lower triangle of the figure indicates the average feature weights of model orders in the row were smaller than the ones in the column, and a positive intensity value indicates the model orders’ average feature weights in the row were larger than the ones in the column. As shows in the figure (Fig. 7), features in the between model orders of 25-50 were consistently more predictive than features in other within and between model orders. Between model order features were consistently more predictive than higher individual model orders (75-100), but less predictive than lower model orders (25-50). And we also see that features in the model order of 50 are more predictive than most other model orders, except the between model order 25-50. Features in the between model 75-100 are consistently less predictive than most within and between model orders, except model of 100.

**Fig. 7.**
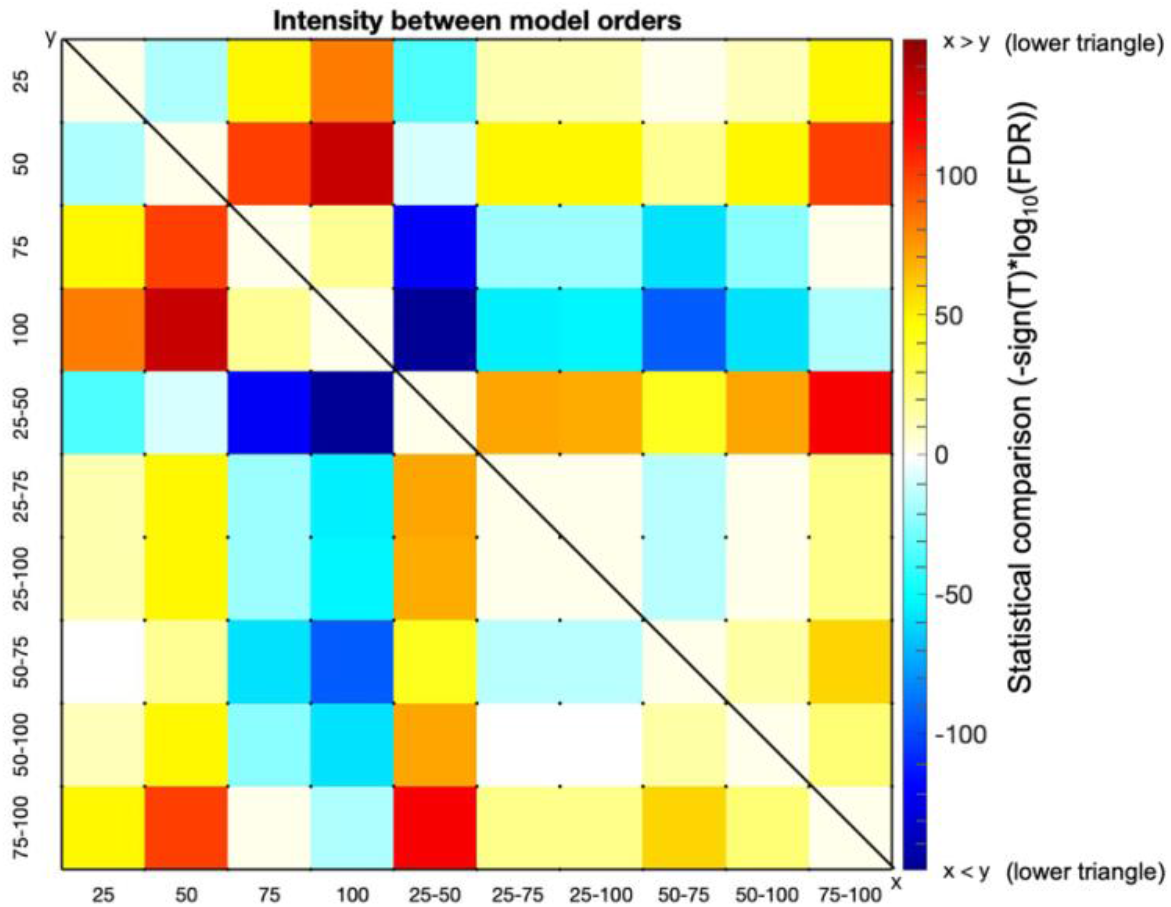
The intensity (-sign(T)^*^log10(FDR)) of average feature weights between model orders. We computed a two-sample t-test between the averaged feature weights of every two model orders, to identify significant differences between them. The negative intensity value in the lower triangle of the figure indicates the average feature weights of model orders in the row (x) were smaller than the ones in the column (y), and the positive intensity value in the lower triangle indicates the model orders’ average feature weights in the row were larger than the ones in the column. The upper triangle shows significant differences (p < 0.05, FDR corrected) between each pair of model orders, whose direction is opposite to the lower triangle.

In order to highlight the variations between domains of model orders, we also calculated the average feature weights (Fig. 8 a.) and maximum feature weights (Fig. 8 b.) in the domain levels. Within each domain, feature weights were aggregated by model orders from 25 to 100. It shows that higher average feature weights were mainly seen in the somatomotor, subcortical, temporal, and visual domains. Strong predictive abilities of FNC features within the somatomotor and visual (highlighted in the figures) were seen at all model orders.

**Fig. 8.**
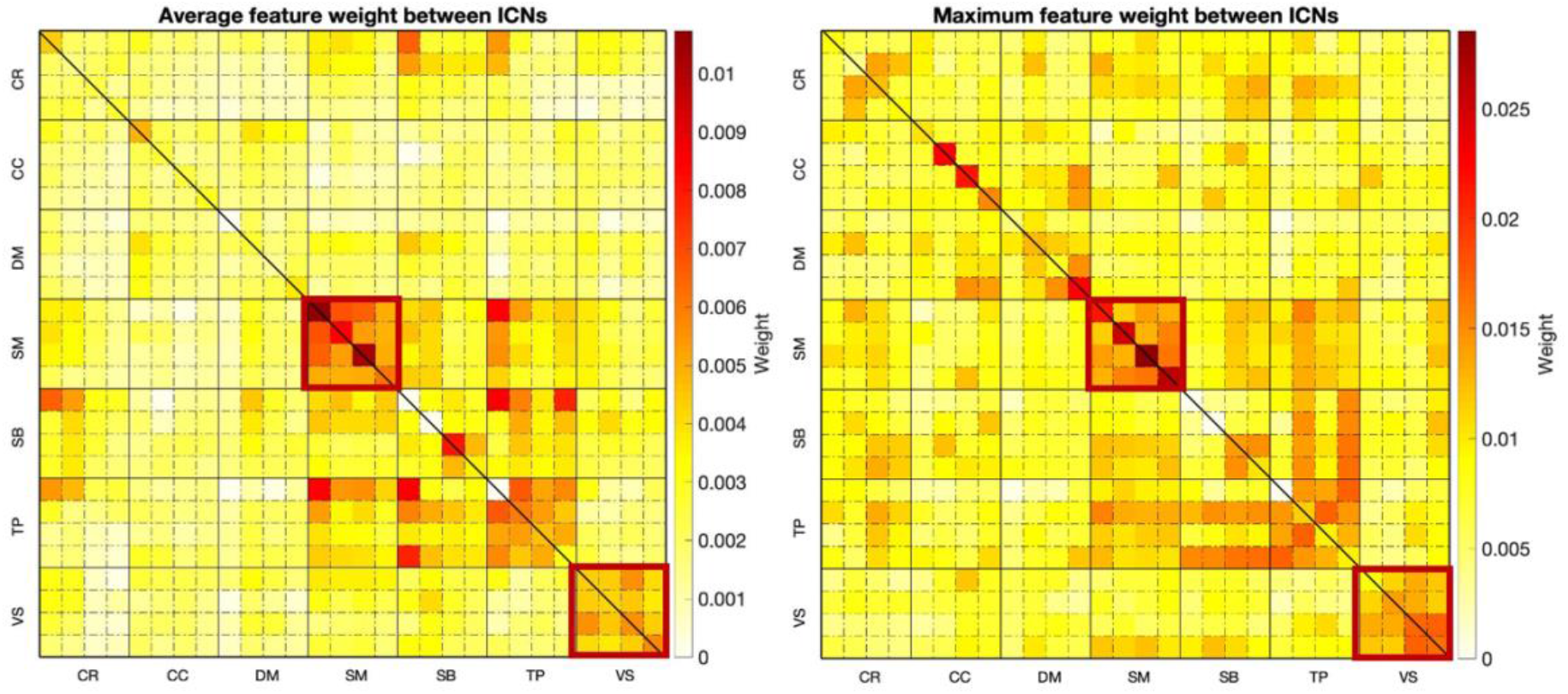
a (left). Average feature weight between domains. Each domain contains four model orders, from 25, 50, 75 to 100, and all the features within each model order were averaged across 100 iterations. Fig. 8 b (right). Maximum feature weight within each domain. Each domain column contains four averaged features of different model orders, from 25, 50, 75 to 100. The maximum feature weight within each domain was selected based on the averaged feature weight across 100 iterations. It shows that higher average feature weights are mainly seen in somatomotor, subcortical, temporal, and visual.

We then performed the two-sample t-test between domains of different model orders(Cressie and Whitford 1986), to identify significant differences between domains. We calculated the intensity values between domains of different model orders (as shown in Fig. 9). For each ICN in the figure, we averaged all the feature weights within each model order of 25, 50, 75, and 100. It indicates that features in somatomotor were always predictive than features in other domains at all model orders, like cerebellum, cognitive control, and default mode (blue regions in the somatomotor row of the lower triangle), and subcortical, temporal, and visual (hot regions in the somatomotor column of the lower triangle). Besides, features in visual were generally predictive than other domains at all model orders, except somatomotor. Features of lower model orders (25-50) in subcortical were generally less predictive than features in other domains.

**Fig. 9.**
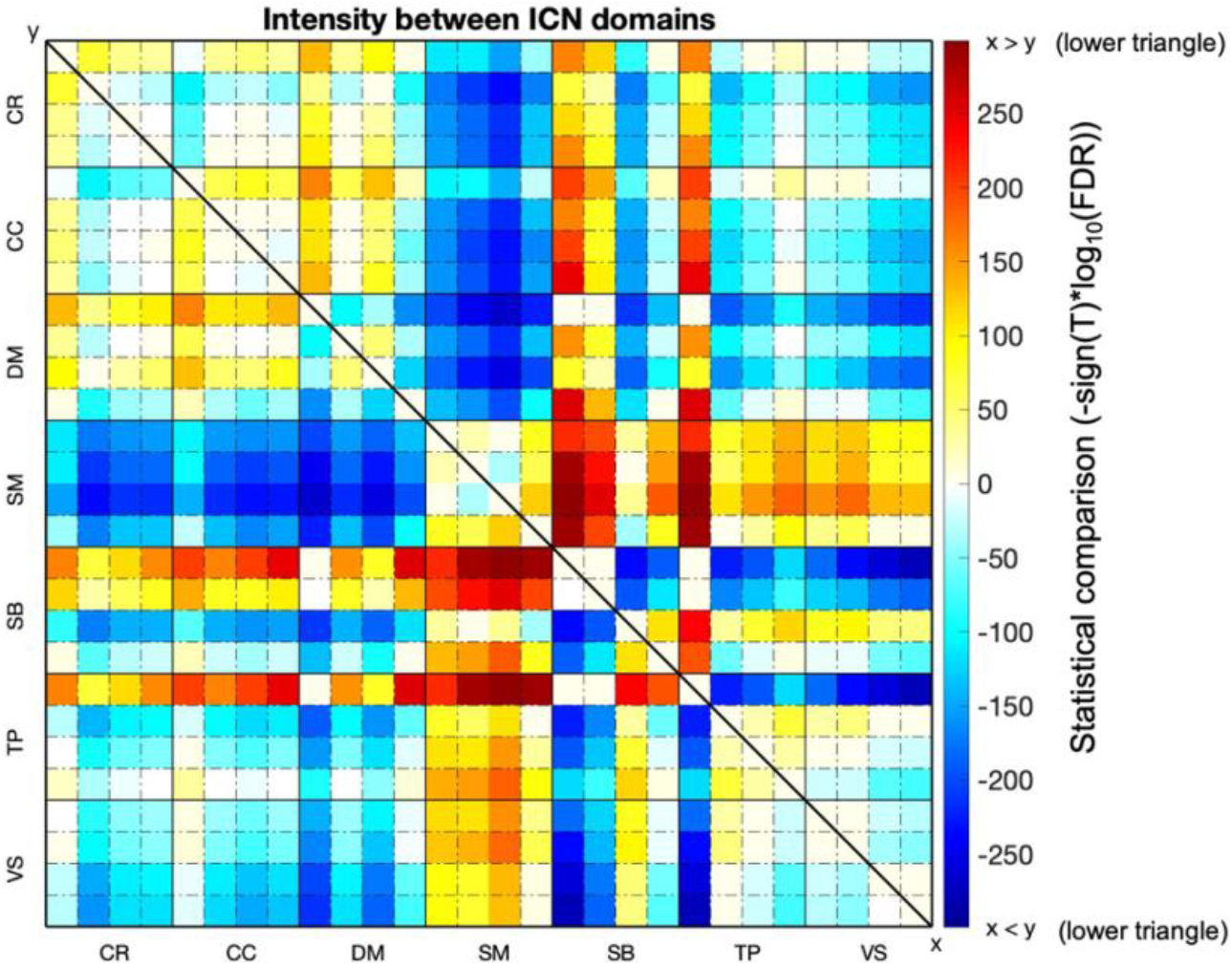
The intensity (-sign(T)^*^log10(FDR)) of average feature weights between domains. The negative intensity values in the lower triangle indicate smaller average feature weights of the ICN in the row (x) compared to the ones in the column (y), and the positive intensity values in the lower triangle indicate larger average feature weights of ICN in the row compared to the column. The upper triangle shows the significant differences (p < 0.05, FDR corrected) between each pair of domains, whose direction is opposite to the lower triangle.

We additionally evaluated the performance of the SVM model. As shown in Table 1, the average accuracy of the SVM model using 100 iterations was 77.4% (outperformed the null accuracy 57.6%), with a specificity of 85.2%, 67.2% sensitivity, and 71.5% F1 score.

**Table 1.** The average performance of SVM model for 100 iterations.

|  | **Accuracy** | **Specificity** | **Sensitivity** | **F1** |
| --- | --- | --- | --- | --- |
| **SVM** | 0.7743 | 0.8522 | 0.6717 | 0.7153 |

Finally, we compared the predictive accuracies of SVM models built by different model orders. We individually trained the SVM model using each number of model order, and then we computed a two-sample t-test between the accuracies of every two SVM models of different model orders. As it shows in the figure (Fig. 10), higher individual model orders always have higher accuracies compared to lower individual model orders. This is due to the fact that including more model orders always resulted in the increase of predictive features. It is also observed that the between model orders of 50-75, 50-100, and 75-100 outperformed individual model orders of 25, 50, 75, and 100, while individual model orders (50-100) outperformed lower between model orders of 25-50 and 25-75.

**Fig. 10.**
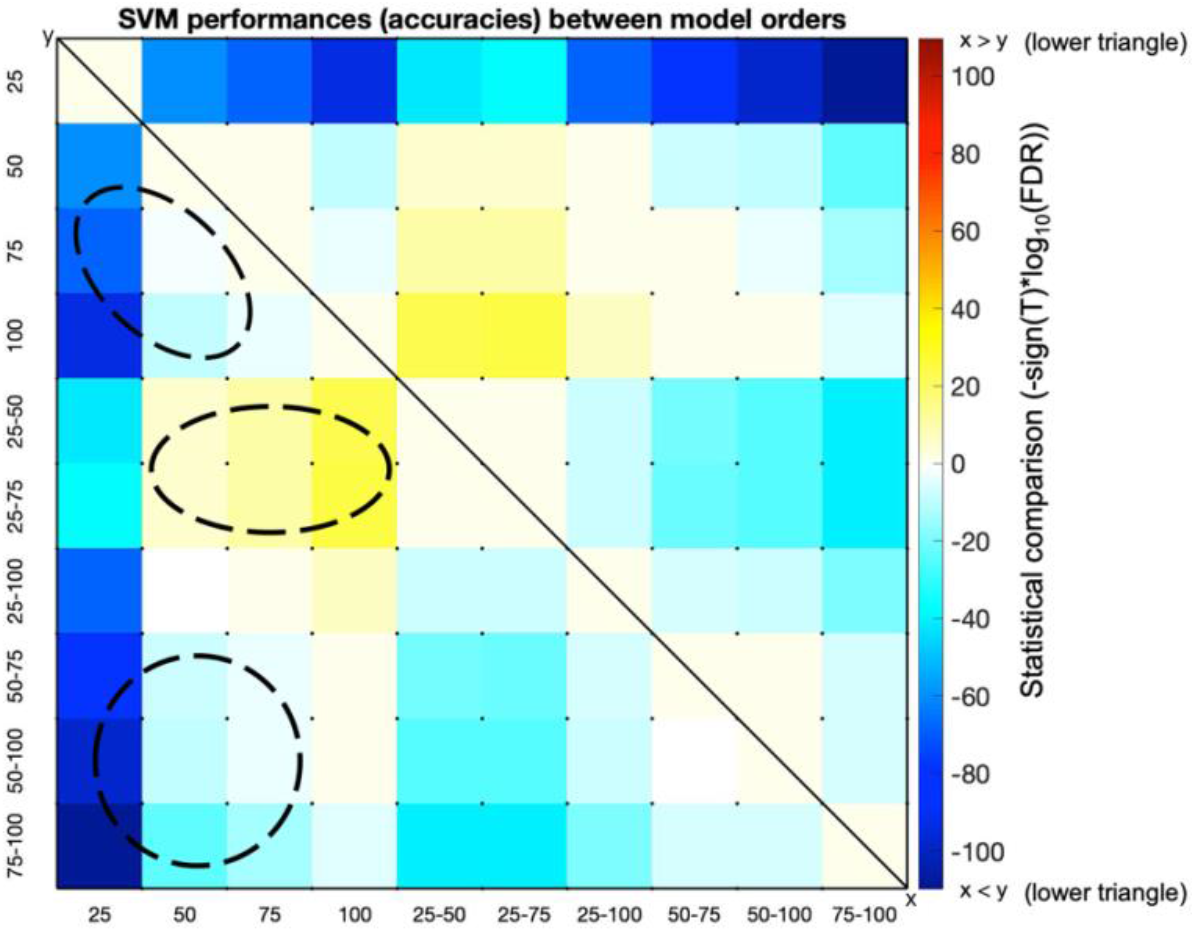
The intensity (-sign(T)^*^log10(FDR)) of SVM accuracies between model orders. The negative intensity value in the lower triangle of the figure indicates the average feature weights of model orders in the row (x) were smaller than the ones in the column (y), and the positive intensity value in the lower triangle indicates the model orders’ average feature weights in the row were larger than the ones in the column. The upper triangle shows the significant differences (p < 0.05, FDR corrected) between each pair of model orders, whose direction is opposite to the lower triangle. The top circle indicates higher individual model orders always have higher accuracies compared to lower individual model orders. The middle circle shows that model orders of 50-100 outperform lower between model orders. The bottom circle indicates higher between model orders outperform individual model orders.

## Discussion

In this paper, we introduce a multi-model order ICA approach to estimate FNCs at multi-spatial scales. We investigated whether multi-spatial-scale FNC features were able to discriminate between schizophrenia individuals and healthy control groups, and found both consistency and uniqueness of FNC patterns within and between model orders. Importantly, we revealed that additional information can be preserved in the between-order FNC that might be ignored in single model order analysis. In addition, some interesting findings were only observed at lower model orders or higher model orders. Specifically, the HC-SZ differences were observed between subcortical vs. temporal and subcortical vs. cerebellum mostly at lower model order FNC (25∼75), but between somatomotor vs. visual networks were mainly at higher model order (100). Results highlighted findings we learned from multiple model orders that would have been missed otherwise.

In addition, we see consistency between the identified significant group differences in FNC and the SVM feature weights, (as shown in Fig. 5 and Fig. 6). We observed that HC generally showed high FNC in somatomotor, visual at all model orders compared to SZ (Fig. 5 a), and these cells were also found to be the most predictive features in the SVM model (Fig. 6 a). For between model orders, we observed that although somatomotor vs. subcortical and visual vs. subcortical were relatively high in SZ compared to HC, they were less predictive than somatomotor and visual. It’s also interesting that some noticeable differences between HC and SZ in the difference mean FNC matrix (Fig. 5 a) do not necessarily lead to their predictive abilities (the same areas in Fig. 6 a are not highlighted). For example, SZs were generally higher than HCs in cerebellum vs. Visual and cerebellum vs. temporal in the difference mean FNC matrix (Fig. 5 a), but they were not noticeably predictive in the SVM model (Fig. 6 a).

Some other useful findings were also indicated by jointly studying the functional brain imaging data within and between the different number of model orders. By comparing the FNC feature predictive ability between different model orders, we surprisingly found that the between model orders always maintained useful information in detecting group differences, particularly, the features between-model order of 25-50 were the most SZ discriminative ones compared to features in other within and between model orders. And in many cases, features in the between model orders were more predictive than individual model order according to our observations, suggesting that investigation of between-model-order FNC may provide informative details that could be missed otherwise. Results also highlighted the differentiating power of somatomotor and visual domain on classifying HC vs. SZ both within and between model orders, compared to other domains.

There are several limitations to this study. First, to obtain a stable feature set, we performed 10 rounds of feature selection within each iteration when turning the SVM model, and within each round of feature selection, we implemented a strategy similar to bootstrap sampling (Breiman 1994), that selecting a subset (50%) of the training data for feature selection. In future work, the performance of the classification model can be improved by increasing the rounds of feature selection and by exploring a range of selected subsets for feature selection, to get a more stable feature set. Second, we selected the top 70% ranked FNC features to build the SVM model, based on our previous experience, that in general, selecting top 70% ranked features maintains the maximum predictive information of the dataset by eliminating redundant features. Going forward we can improve the performance of the classification model by exploring a range of selected features, to get an optimal predictive model. We tried both using feature selection and not using it to build the SVM model, the results showed that the performance of using feature selection was slightly better than not using it. In addition to the SVM model, we also tried another classification model random forest (Ho 1995), the performance of SVM was slightly better compared to the random forest for our case, so we keep SVM results. As future work, more efforts can be made to explore other machine learning techniques, like boosting (Schapire 2003), to improve the accuracy of the learning classifier.

## Conclusions

In summary, based on fMRI data from 477 healthy participants and 350 schizophrenia patients, we compared the group differences obtained at different model orders by evaluating the multi-order FNCs, as well as testing the classification power of FNC at different model orders. A comprehensive visualization of the relationships via FNC block plots and FNC finger plots was introduced. Results suggest that additional information was observed both at different model orders and in between model order relationships, which goes beyond the known general modular relationships in ICNs. We are also releasing four model-order templates to the public for future use. Our study highlights the benefit of studying functional network connectivity within and between multiple spatial scales. This work expands upon and adds to the relatively new literature on resting fMRI-based classification and prediction.

## Acknowledgments

Thanks to the studies which collected the original data.

## Authorship Confirmation Statement

Xing Meng: Conceptualization; Formal analysis; Investigation; Methodology; Visualization; Writing – original draft; Writing – review & editing. Armin Iraji: Conceptualization; Investigation; Methodology; Visualization; Writing – review & editing. Zening Fu: Data collection and preprocessing. Peter Kochunov: Data collection; Resources. Aysenil Belger: Data collection; Resources. Judy Ford: Data collection; Resources. Sarah McEwen: Data collection; Resources. Daniel Mathalon: Data collection; Resources. Bryon Mueller: Data collection; Resources. Godfrey Pearlson: Data collection; Resources. Steven G. Potkin: Data collection; Resources. Adrian Preda: Data collection; Resources. Jessica Turner: Data collection; Resources. Theodorus Van Erp: Data collection; Resources. Jing Sui: Writing – review & editing. Vince Calhoun: Conceptualization; Funding acquisition; Investigation; Methodology; Project administration; Resources; Writing – review & editing.

